# SPLIT HAND/FOOT VARIANTS FAIL TO RESCUE PRDM1A MUTANT CRANIOFACIAL DEFECTS

**DOI:** 10.1101/2023.05.19.541469

**Authors:** Brittany T. Truong, Lomeli C. Shull, Ezra Lencer, Kristin B. Artinger

**Affiliations:** Human Medical Genetics & Genomics Graduate Program, University of Colorado Denver Anschutz Medical Campus, Aurora, CO, 80045, USA; Department of Craniofacial Development, University of Colorado Denver Anschutz Medical Campus, Aurora, CO, 80045, USA; Biology Department, Lafayette College, Easton, PA, 18042, USA; Department of Diagnostic and Biological Sciences, University of Minnesota School of Dentistry, Minneapolis, MN, 55455, USA

**Author notes:** **Correspondence:** Kristin Bruk Artinger, Diagnostic and Biological Sciences, University of Minnesota School of Dentistry, 515 Delaware St SE, 28 Minneapolis, MN, 55455 **Email:**.

**Keywords:** PRDM1, Split Hand/Foot Malformation, craniofacial, zebrafish

## Abstract

**Background:** Split Hand/Foot Malformation (SHFM) is a congenital limb disorder presenting with limb anomalies, such as missing, hypoplastic, or fused digits, and often craniofacial defects, including a cleft lip/palate, microdontia, micrognathia, or maxillary hypoplasia. We previously identified three novel variants in the transcription factor, *PRDM1*, that are associated with SHFM phenotypes. One individual also presented with a high arch palate. Studies in vertebrates indicate that PRDM1 is important for development of the skull; however, prior to our study, human variants in *PRDM1* had not been associated with craniofacial anomalies.

**Methods:** Using transient mRNA overexpression assays in *prdm1a*^*-/-*^ mutant zebrafish, we tested whether the *PRDM1* SHFM variants were functional and could lead to a rescue of the craniofacial defects observed in *prdm1a*^*-/-*^ mutants. We also mined a CUT&RUN and RNA-seq dataset to examine Prdm1a binding and the effect of Prdm1a loss on craniofacial genes.

**Results:** *prdm1a*^*-/-*^ mutants exhibit craniofacial defects including a hypoplastic neurocranium, a loss of posterior ceratobranchial arches, a shorter palatoquadrate, and an inverted ceratohyal. Injection of wildtype *hPRDM1* in *prdm1a*^*-/-*^ mutants partially rescues these structures. However, injection of each of the three SHFM variants fails to rescue the skeletal defects. Loss of *prdm1a* leads to a decreased expression of important craniofacial genes, such as *dlx5a/dlx6a, hand2, sox9b, col2a1a*, and *hoxb* genes.

**Conclusion:** These data suggest that the three SHFM variants are not functional and may have led to the craniofacial defects observed in the humans. Finally, they demonstrate how Prdm1a can directly bind and regulate craniofacial gene expression.

## Introduction

Split Hand/Foot Malformation (SHFM) is a congenital limb disorder affecting 1 in 18,000 live births (Umair & Hayat, 2020). Individuals with SHFM present with missing, hypoplastic, and/or fused digits, though the phenotypes are highly variable due to incomplete penetrance. Many individuals also present with craniofacial defects, such as a cleft lip/palate, microdontia, micrognathia, and/or maxillary hypoplasia (reviewed in Sowinska-Seidler et al. (2014)). Pathogenic variants in *WNT10B* (MIM #225300), *TP63* (MIM #605289), *DLX5* (MIM #183600), *ZAK* (MIM #616890), *EPS15L1* (MIM *616826), or chromosomal rearrangements in chromosomes 2 (MIM %606708), 10 (MIM #246560), or X (MIM %313350) are known to cause SHFM (Sowinska-Seidler et al., 2014). We recently identified three novel, pathogenic variants in a transcription factor, *PRDM1*, in individuals with SHFM: *PRDM1c*.*712_713insT* (p.C239Lfs*32); *PRDM1c*.*1571C>G* (p.T524R); and *PRDM1c*.*2455A>G* (p.T819A) (Truong et al., 2023). These variants negatively affect the protein’s ability to bind to DNA and regulate genes required for limb induction, outgrowth, differentiation, and anterior/posterior patterning (Truong et al., 2023). One individual also has a craniofacial defect. He came into the clinic with ectrodactyly ectodermal dysplasia (EEC) syndrome (MIM 129810) with bilateral 3/4-digit syndactyly and a high arch palate but no clefting. This individual has a *de novo*, missense mutation, *PRDM1c*.*1571C>G* (p.T524R).

Studies in zebrafish and mice indicate that PRDM1 is important for development of the skull. Much of the craniofacial skeleton is derived from neural crest cells (NCCs), a multipotent population of cells derived from the non-neural ectoderm at the neural plate border. During development, these cells migrate away from the neural tube into different regions of the body and differentiate into various cell types, such as peripheral neurons, osteoblasts, melanocytes, and chondrocytes. Cranial NCCs migrate into the pharyngeal arches and facial prominences of the developing face. These cells will later contribute to the bone, cartilage, and connective tissue of the developing head skeleton.

In zebrafish, Prdm1a has been shown to be required for NCC specification and differentiation, as indicated by a significant decrease in NCC markers *sox10, crestin*, and *snail2* in *prdm1a*^*-/-*^ presumed null and hypomorph mutants (Artinger et al., 1999; Hernandez-Lagunas et al., 2005; Olesnicky et al., 2010; Roy & Ng, 2004). In turn, a loss of Prdm1a leads to a decrease in NCC derivatives, including melanocytes and cranial and dorsal root ganglia (Artinger et al., 1999; Birkholz et al., 2009; Hernandez-Lagunas et al., 2005; Olesnicky et al., 2010; Roy & Ng, 2004). *prdm1a*^*-/-*^ mutants present with craniofacial defects, including an inverted ceratohyal, missing posterior ceratobranchial arches, and a shortened neurocranium (Birkholz et al., 2009). In mice, *Prdm1* is expressed in the endodermal layer of the first branchial arch at E9.5 and in the endoderm, ectoderm, and mesenchyme of the second and third arches (Robertson et al., 2007; Vincent et al., 2005). In both null *Prdm1* mice and conditional *Prdm1* knockouts in the embryo proper (*Sox2:Cre*), mutant embryos are able to properly form the first pharyngeal arch, and subsequently a lower jaw, but the more caudal arches are completely lost. Mutants are missing the thymus and exhibit hypoplasia of the pharyngeal epithelium (Robertson et al., 2007; Vincent et al., 2005). These studies provide evidence for the importance of PRDM1 in craniofacial development.

Prior to our study, human variants in *PRDM1* had not been associated with craniofacial anomalies. Using transient overexpression assays, we sought to determine whether the *PRDM1* SHFM variants were functional and could rescue the craniofacial defects in *prdm1a*^*-/-*^ zebrafish.

## Results

### SHFM Human *PRDM1* variants lead to craniofacial defects

To determine whether the SHFM *hPRDM1* variants are functional, we designed an *in vivo* rescue experiment. We overexpressed either wildtype *hPRDM1* mRNA or each of the three SHFM variants in intercrossed *prdm1a*^*+/-*^ zebrafish embryos and assessed whether there was a rescue to the craniofacial skeleton (**Fig. 1**). Uninjected *prdm1a*^*-/-*^ mutants are missing posterior ceratobranchial arches, have an inverted and hypoplastic ceratohyal (16.28% decrease, p=0.0125), shorter palatoquadrate (21.57% decrease, p=0.0079), narrow ethmoid plate (14.46% decrease, p=0.0780), and a shortened neurocranium (16.56% decrease, p=0.0397) compared to uninjected wildtype/heterozygotes (**Fig. 1A-B, H-K**). Injection of wildtype *hPRDM1* partially rescues the head, particularly the length of the palatoquadrate (24.25% increase, p=0.0256) (**Fig. 1C, I**). However, injection of the three SHFM variants fails to rescue the craniofacial skeleton (**Fig. 1C-F**). The posterior ceratobranchial arches are consistently missing (**Fig. 1C-G**), and the ceratohyal, palatoquadrate, and ethmoid plate remained shorter (**Fig. 1H-K**). There is a slight increase in the length of the neurocranium from the ethmoid plate to the trabeculae upon injection with wildtype *hPRDM1, PRDM1* (p.T524R), and *PRDM1* (p.T819A), but the results were not statistically significant (**Fig. 1J**). Inability of the wildtype allele to fully rescue is likely due to the transience of the assay, insufficient dosing and/or rapid degradation of the mRNA. Together, these data suggest that the SHFM *PRDM1* variants are not functional and may lead to craniofacial defects.

**Figure 1:**
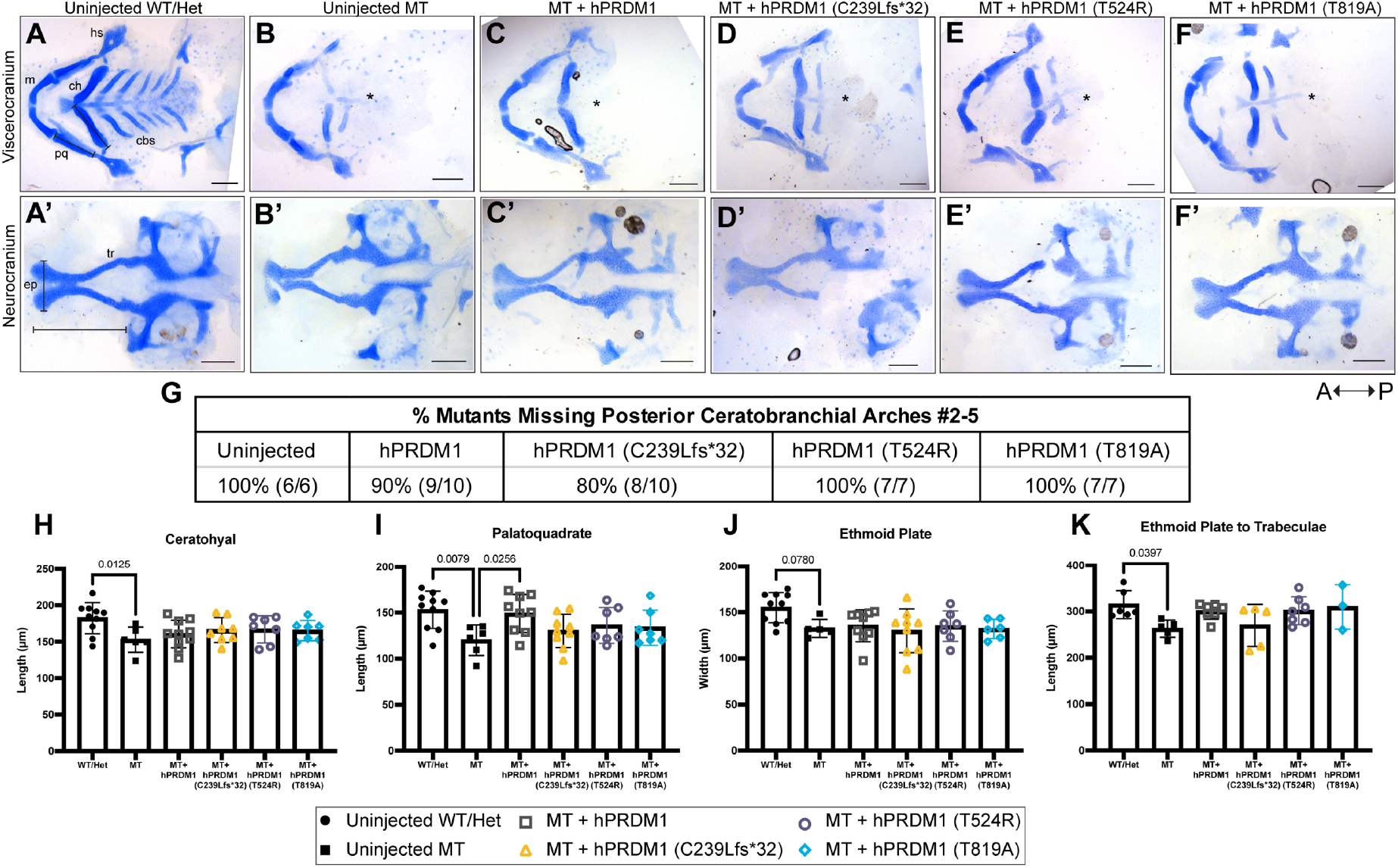
Transient overexpression of wildtype *hPRDM1* in *prdm1a*^*-/-*^ embryos rescues palatoquadrate, while the SHFM variants fail to rescue. *prdm1a*^*+/-*^ heterozygous fish were intercrossed and injected with the *hPRDM1* wildtype and SHFM variant mRNA at the single-cell stage. Injected larvae were collected at 4 dpf for Alcian blue staining. (**A-F**) Representative images of dissected craniofacial skeletons. The top row contains dissections of the viscerocranium, while the bottom row is the neurocranium. (**A**) Uninjected wildtype/heterozygous (n=10). (**B**) Uninjected *prdm1a*^*-/-*^ mutant (n=6). *prdm1a*^*-/-*^ mutants were injected with (**C**) wildtype *hPRDM1* (n=10), (**D**) *hPRDM1*(p.C239Lfs*32) (n=10), (**E**) *hPRDM1*(p.T524R) (n=7), or (**F**) *hPRDM1*(p.T819A) mRNA (n=7). The asterisk (*) represents missing ceratobranchial arches. (**G**) Table showing the proportion of *prdm1a-/-* mutants that had missing posterior ceratobranchial arches #2-5. Measurements were taken to quantify the (**H**) length of the ceratohyal, (**I**) length of the palatoquadrate, (**J**) width of the ethmoid plate, and (**K**) length from the ethmoid plate to trabeculae. *prdm1a*^*-/-*^ mutants have missing posterior ceratobranchial arches and a shorter ceratohyal, palatoquadrate, ethmoid plate, and neurocranium overall. Injection of wildtype *hPRDM1* rescues the length of the palatoquadrate (p=0.0256), while the other variants do not. Injection of wildtype *hPRDM1, hPRDM1*(p.T524R), and *hPRDM1*(p.T819A) modestly rescues the length of the neurocranium. Abbreviations: cbs, ceratobranchial arches; ep, ethmoid plate; hs, hyosymplectic; m, Meckel’s cartilage; pq, palatoquadrate; tr, trabeculae

### Loss of Prdm1a leads to decreased expression of craniofacial genes

PRDM1 has been shown to be involved in vertebrate craniofacial development, though the mechanistic role remains unclear (Birkholz et al., 2009; Robertson et al., 2007; Vincent et al., 2005). We examined an RNA-seq dataset from isolated pectoral fin cells at 48 hours post fertilization (hpf) using a *Tg(Mmu:Prx1-EGFP)* transgenic line that labels the pectoral fin, pharyngeal arches, and dorsal part of the head (Truong et al., 2023). Though we dissected and removed the head prior to sorting the cells, there was some arch and dorsal head tissue in the sample. In turn, one of the most significant downregulated genes in *prdm1a*^*-/-*^ mutants was *emilin3a*, a glycoprotein within the extracellular matrix belonging to the EMILIN/multimerin family (**Fig. 2A**). This gene is expressed in the notochord, pharyngeal arches, and developing craniofacial skeleton of zebrafish, but not in the pectoral fin or limb (Milanetto et al., 2007); however, its role has not yet been explored. It is possible that Prdm1a and Emilin3a interact during craniofacial development. Additional genes that were downregulated and known to be expressed in the pharyngeal arches or are involved in craniofacial development include *sox10, hand2, dlx2a, dlx5a/dlx6a, gsc, barx1, prdm3, sox9b, col2a1a*, and members of the *hoxb* gene family (**Fig. 2A**) (Dougherty et al., 2012; Miller et al., 2003; Robledo et al., 2002; Shull et al., 2020; Sperber et al., 2008; Yamada et al., 2021; Yan et al., 2005).

**Figure 2:**
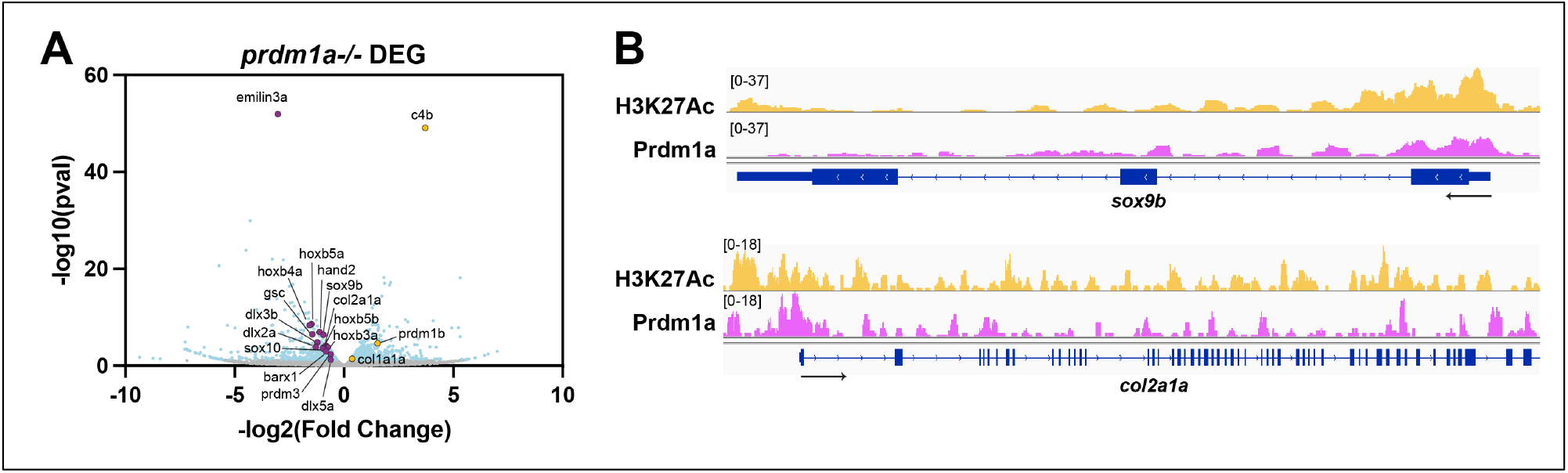
Loss of Prdm1a leads to a significant decrease in genes required for craniofacial development. (**A**) RNA-seq was performed on *Tg(Mmu:Prx1-EGFP)* wildtype and *prdm1a*^*-/-*^ embryos at 48 hpf. Volcano plot showing spread of differentially genes in pectoral fins of *prdm1a*^*-/-*^ compared to wildtype. Light blue dots are significant, differentially expressed genes (-log_10_ p-value?1.15). Purple dots are downregulated, selected genes of interest, while yellow dots are upregulated. (**B**) CUT&RUN was performed on *Tg(Mmu:Prx1-EGFP)* wildtype embryos at 24 hpf. Tracks showing H3K27Ac enrichment (open chromatin) and Prdm1a binding sites for *sox9b* and *c****o****l2a1a*.

We also examined a CUT&RUN dataset on the *Tg(Mmu:Prx1-EGFP)* transgenic line at 24 hpf to test direct Prdm1a binding to DNA. We mined the data and found that Prdm1a directly binds putative enhancer and promoter regions upstream of *sox9b, col2a1a*, and *dlx5a/dlx6a* (**Fig. 2B**) (Truong et al., 2023). It is possible that Prdm1a helps regulate cartilage formation in the craniofacial skeleton. Together, our data suggest that Prdm1a is required during craniofacial development.

## Discussion

Although it is typically perceived as a congenital limb disorder, SHFM individuals have been reported to exhibit craniofacial anomalies, including cleft lip/palate, microdontia, micrognathia, and/or maxillary hypoplasia (reviewed in Sowinska-Seidler et al. (2014)). We recently identified three novel, causal SHFM variants in a gene *PRDM1*, which led to clefted hands/feet with variable penetrance (Truong et al., 2023). One individual also presented with a craniofacial defect, namely a high arched palate. We found that the *PRDM1* variants have a dominant negative effect on the developing limb bud and fail to rescue the pectoral fin in *prdm1a*^*-/-*^ mutant zebrafish, providing evidence for their pathogenicity (Truong et al., 2023). Here, we show that the each of the three variants also fails to rescue the craniofacial skeleton, suggesting that the variants are not functional. In contrast, overexpression of wildtype *hPRDM1* mRNA partially restores the face, particularly the length of the palatoquadrate. This was expected given that overexpression of zebrafish *prdm1a* mRNA sufficiently rescues NCCs in mutants (Hernandez-Lagunas et al., 2005). Inability of the wildtype allele to rescue other elements of the craniofacial skeleton, such as the posterior ceratobranchial arches or hypoplastic neurocranium, may be because of the transient nature of the experiment or due to an insufficient dosage of *PRDM1*. Indeed, *prdm1a*^*-/-*^ mutants have severe craniofacial phenotypes with 100% penetrance (**Fig. 1A, B, G**) and may require a higher dose of mRNA in order to recover other structures of the head. Increasing the dose or using a genetically stable approach to overexpress the variants will be useful for better interpreting their function. We also mined a RNA-seq data comparing wildtype and *prdm1a*^*-/-*^ mutants at 48 hpf and found a significant decrease in genes critical for craniofacial development, including *dlx5a/dlx6a, hand2, barx1, gsc, sox9b, col2a1a*, and members of the *hoxb* gene family (**Fig. 2A**) (Truong et al., 2023). Our CUT&RUN data at 24 hpf also suggests that Prdm1a is directly binding to putative enhancers and promoters of *dlx5a/dlx6a, sox9b*, and *col2a1a* (**Fig. 2B**) (Truong et al., 2023).

Of particular interest is *dlx5a/dlx6a*, a set of homeotic genes required for both limb development and dorsal/ventral patterning in the face (Robledo et al., 2002). Variants in *DLX5* are known to cause SHFM Type I (MIM #183600) with orofacial clefting (Bernardini et al., 2008; Elliott & Evans, 2006), and we have shown that Prdm1a directly binds to and regulates *dlx5a* in the pectoral fin (Truong et al., 2023). We hypothesize that Prdm1a is required for maintaining proper patterning in the face through its regulation of *dlx5a/dlx6a* in the pharyngeal arches as well, though additional experiments are needed to determine this. Structures in the craniofacial and limb skeleton are clearly distinct from one another, but they utilize many of the same gene regulatory networks and mechanisms for their development (Truong & Artinger, 2021). This then leads to a frequent co-occurrence of craniofacial and limb anomalies in congenital diseases. These future studies will provide critical insight into the mechanism by which Prdm1a regulates both craniofacial and limb development and how disruptions to the gene regulatory networks involved can lead to SHFM.

## Materials & Methods

### Zebrafish husbandry

Zebrafish were maintained as previously described (Westerfield, 2000). The wildtype (WT) strain used was AB (ZIRC) and the mutant lines used were *prdm1a*^*m805*^ *(nrd;* referred to as *prdm1a*^*-/-*^) (Artinger et al., 1999; Hernandez-Lagunas et al., 2005). All experiments were reviewed and approved by the Institutional Animal Care and Use Committee (IACUC) at the University of Colorado Denver Anschutz Medical Campus (protocol #147) and conform to the NIH regulatory standards of care and treatment. Zebrafish lines can be obtained from the lead contact.

### mRNA overexpression in zebrafish

mRNA overexpression was performed as previously described (Truong et al., 2023). *hPRDM1* variant cDNA was synthesized into a pCS2+ backbone using Gateway cloning. cDNA was linearized and transcribed using the mMessage mMachine T7 Transcription Kit (ThermoFisher). *prdm1a*^*+/-*^ fish were intercrossed and the different *hPRDM1* mRNA variants (diluted 1:10 in water and phenol red) were injected into resulting embryos at the single-cell stage. Embryos were staged throughout the first four days of development to ensure there was no developmental delay or unassociated pathologies due to the mRNA overexpression. At 4 dpf, larvae were collected for Alcian blue staining. Sample size refers to the number of individuals and is included in the figure legends.

### Alcian blue cartilage staining

Zebrafish were stained for cartilage as previously described (Walker & Kimmel, 2007). In short, 4 dpf larvae were fixed in 2% paraformaldehyde (PFA) at room temperature for one hour. Larvae were then washed in 100mM Tris (pH 7.5)/10mM MgCl_2_ before rocking overnight at room temperature in Alcian blue stain (pH 7.5) (0.04% Alcian Blue, 80% ethanol, 100mM Tris [pH 7.5], 10mM MgCl_2_). Larvae were destained and rehydrated in a series of ethanol washes (80%, 50%, 25%) containing 100mM Tris (pH 7.5) and 10mM MgCl_2_, and then bleached for 10 minutes in 3% H_2_O_2_/0.5% KOH. Finally, larvae were rinsed twice in 25% glycerol/0.1% KOH to remove the bleach and stored at 4°C in 50% glycerol/0.1% KOH. The craniofacial skeletons of stained larvae were dissected, flat mounted in 50% glycerol/0.1 KOH, and imaged on an Olympus BX51 WI microscope. Measurements of the viscerocranium and neurocranium were performed blindly in ImageJ and then compared using a one-way ANOVA followed by a Tukey’s post-hoc test relative to uninjected *prdm1a*^*-/-*^ mutants. Sample size refers to the number of individuals and is included in the figure legends.

### Fluorescence-activated cell sorting (FAC)

EGFP-positive cells were isolated from zebrafish embryos using fluorescence-activated cell (FAC) sorting as previously described (Truong et al., 2023). In short, *Tg(MmuPrx1:EGFP)* embryos were collected at 24 hpf (CUT&RUN) and 48 hpf (RNA-seq). Embryos were pooled together, washed in DPBS (Gibco), and dissociated in Accumax (Innovative Cell Technologies) and DNase I (50 units/100 embryos) (Roche) at 31°C for 1.5 hours. Cells were homogenized and washed in solution (300 units DNase I in 4 mL DPBS) before filtering through a 70 μM nylon mesh cell strainer (Fisher Scientific). Cells were spun down, resuspended in basic sorting buffer (1mM EDTA, 25mM HEPES [pH 7.0], 1% FBS in DPBS), stained with DAPI (1:1000), and FAC sorted at the University of Colorado Cancer Center Flow Cytometry Shared Resource (Aurora, CO, USA) on the MoFlo XDP100 sorter (Beckman Coulter) with a 100 μM nozzle tip.

### CUT&RUN

Following the FAC sort, cleave under targets and release using nuclease (CUT&RUN) was performed on 150,000+ sorted EGFP-positive pectoral fin cells at 24 hpf as previously described (Truong et al., 2023). Briefly, cells were incubated on activated Concanavalin A conjugated paramagnetic beads (EpiCypher) at room temperature for 10 minutes. Cells were washed in Antibody Buffer (20mM HEPES, pH 7.5; 150 mM NaCl; 0.5 mM Spermidine [Invitrogen], 1X Complete-Mini Protease Inhibitor tablet [Roche Diagnostics]; 0.01% Digitonin [Sigma-Aldrich]; 2 mM EDTA) and incubated overnight at 4°C with rotation in the respective antibody (IgG [1:100; Jackson ImmunoResearch, 111-005-003, RRID: AB_2337913], H3K27ac [1:66; Cell Signaling Technology, 4353S, RRID: AB10545273], and Prdm1a [1:33; rabbit polyclonal antibody from Dr. Phillip Ingham (von Hofsten et al., 2008); validated for chromatin immunoprecipitation (ChIP) by Powell et al. (2013)]. Excess antibody was removed by washing in cold Digitonin Buffer (20 mM HEPES, pH 7.5; 150 mM NaCl; 0.5 mM Spermidine; 1X Complete-Mini Protease Inhibitor tablet). Cells were then incubated with pAG-MNase (EpiCypher) for 10 minutes at room temperature and washed with Digitonin Buffer. Cells were rotated in 100 mM CaCl_2_ at 4°C for two hours before the Stop Buffer (340 mM NaCl, 20 mM EDTA, 4 mM EGTA, 50 μg/ml RNaseA, 50 μg/ml glycogen) was added for 10 minutes at 37°C without the *E*.*coli* spike-in. DNA fragments were purified with a DNA Clean & Concentrate Kit (Zymo Research). Eluted DNA fragments were amplified using the NEBNext Ultra II DNA Library Prep Kit for Illumina (New England Biolabs) following the manufacturer’s instructions. Amplification of DNA was performed following guidelines outlined by EpiCypher: 98°C 45s, [98°C 15s, 60°C 10s] x14 cycles, 72°C 1 min. Samples were subjected to paired end 150 bp sequencing on the Illumina NovaSEQ 6000 system at Novogene Corporation Inc. (Sacramento, CA). CUT&RUN experiments were performed in duplicate for two biological replicates. Bioinformatics analysis was performed as previously described (Truong et al., 2023).

### RNA-sequencing

RNA-sequencing was performed as previously described (Truong et al., 2023).About 250 *Tg(MmuPrx1:EGFP);prdm1a*^*-/-*^ (sorted by pigment phenotype (Artinger et al., 1999; Hernandez-Lagunas et al., 2005)) and *Tg(MmuPrx1:EGFP)* wildtype embryos were dissected at 48 hpf to remove the brain before FAC sorting. RNA from sorted cells was extracted using the RNAqueous™-Micro Total RNA Isolation Kit (ThermoFisher) following the manufacturer’s instructions for cultured cells. DNase treatment was performed. A library was prepared using the NEBNext Ultra II Directional RNA Library Kit for Illumina (New England Biolabs) following the manufacturer’s instructions. Samples were subjected to sequencing on the Illumina NovaSEQ 6000 system at Novogene Corporation Inc. (Sacramento, CA) at a depth of over 20 million reads per sample. RNA-seq experiments were performed in duplicate for two biological replicates per genotype. Bioinformatics analysis was performed as previously described (Truong et al., 2023).

## Data Availability

The RNA-seq and CUT&RUN data have been deposited in NCBI’s Gene Expression Omnibus and are accessible through accession number GSE217486 (https://www.ncbi.nlm.nih.gov/geo/query/acc.cgi?acc=GSE217486).

## Acknowledgements

We thank members of the Artinger Lab for project feedback; Christine Archer and the zebrafish facility team for excellent animal care; and the SHFM families for participation in the study.

## Author Contributions

Conceptualization: K.B.A.; Methodology: B.T.T., L.C.S.; Formal analysis: B.T.T., L.C.S., E.L.; Writing – original draft preparation: B.T.T., K.B.A.; Writing – review and editing: L.C.S., E. L.; Supervision: K.B.A.; Funding acquisition: B.T.T., K.B.A.

